# HGDiscovery: an online tool providing functional and phenotypic information on novel variants of homogentisate 1,2- dioxigenase

**DOI:** 10.1101/2021.04.26.441386

**Authors:** Malancha Karmakar, Vittoria Cicaloni, Carlos H.M. Rodrigues, Ottavia Spiga, Annalisa Santucci, David B. Ascher

**Affiliations:** Computational Biology and Clinical Informatics, Baker Heart and Diabetes Institute, Melbourne, Victoria, Australia; Structural Biology and Bioinformatics, Department of Biochemistry, University of Melbourne, Melbourne, Victoria, Australia; Systems and Computational Biology, Bio21 Institute, University of Melbourne, Melbourne, Victoria, Australia; Department of Biotechnology, Chemistry and Pharmacy, University of Siena, Siena, Italy; Department of Biochemistry, Bio21 Institute, University of Cambridge, Cambridge, UK

## Abstract

Alkaptonuria (AKU), a rare genetic disorder, is characterized by the accumulation of homogentisic acid (HGA) in the body. Affected individuals lack enough functional levels of an enzyme required to breakdown HGA. Mutations in the *HGD* gene cause AKU and they are responsible for deficient levels of functional homogentisate 1,2-dioxygenase (HGD), which, in turn, leads to excess levels of HGA. Although HGA is rapidly cleared from the body by the kidneys, in the long term it starts accumulating in various tissues, especially cartilage. Over time (rarely before adulthood), it eventually changes the color of affected tissue to slate blue or black. Here we report a comprehensive mutation analysis of 111 pathogenic and 190 non-pathogenic HGD missense mutations using protein structural information. Using our comprehensive suite of graph-based signature methods, mCSM complemented with sequence-based tools, we studied the functional and molecular consequences of each mutation on protein stability, interaction and evolutionary conservation. The scores generated from the structure and sequence-based tools were used to train a supervised machine learning algorithm with 84% accuracy. The empirical classifier was used to generate the variant phenotype for novel HGD missense mutations. All this information is deployed as a user friendly freely available web server called HGDiscovery (http://biosig.unimelb.edu.au/hgdiscovery/).

## Introduction

Alkaptonuria (AKU) is a rare recessive metabolic disorder which was used by Sir Archibald Garrod in his Croonian lectures to describe inborn errors of metabolism [1]. It is a hereditary disorder, resulting from mutations in the enzyme homogentisate 1,2 dioxygenase (HGD) (EC 1.13.11.5), responsible for the breakdown of homogentisic acid (HGA) which is an intermediate metabolite in the tyrosine degradation pathway [2]. With blockage in tyrosine metabolism, elevated levels of HGA leads to deposition of its own polymers as an ochronotic pigment in the connective tissue including cartilage, heart valves, and sclera [3]. Manifestation of disease during early childhood is seen as “homogentisic aciduria”, which is darkening of the urine upon standing. Delayed symptoms can be seen after 30 years of age which involves “ochronosis” – pigmentation of collagenous tissues like cardiac valves, eyes, ears and skin [4]. Current estimates of the disease occurrence in the Unites States obtained from the National Organisation of Rare Disorders is 1 in 250,000 – 1,00,000 live births [5].

HGD gene located on chromosome 3q21-q23 [6], is a single copy gene composed of 14 exons [7]. Due to compound heterozygosity or homozygosity of HGD gene variants, the enzymatic defect in HGD is autosomal recessive [6, 8]. Information on all variants identified till date globally have been documented in the HGD mutation database (http://hgddatabase.cvtisr.sk/). The experimental crystal structure of the HGD protein has been solved (PDB code 1EY2 and 1EYB) in 2000. The HGD protein protomer (NP_000178.2), is composed of 445 amino acids, which includes a 280 residue N-terminal domain, a central β-sandwich and a 140 residue C-terminal domain [8]. It is a complex hexameric protein arranged as a dimer of trimers [9]. It is principally expressed in osteoarticular compartment cells (i.e. chondrocytes, synoviocytes and osteoblasts) [10] and in prostate, small intestine, colon, kidney and liver [7]. The spatial structure of the protomer, two-disc like trimers and the hexamer are maintained by an intricate network of non-covalent inter and intra-molecular interaction. This makes the protein structure extremely vulnerable to mutations [11].

The major obstacle in studying an ultra-rare and complex disease like AKU is the lack of a standardized methodology to assess disease severity and response to treatment [12], which is complicated by the fact that AKU symptoms differ from one individual to another. Detailed evaluation and comparison of clinical and genomic data of AKU patient can play a key role to understand AKU variability. An in-depth molecular characterization of the disease is needed in pharmacogenomics prediction for suitable medical treatment. To address the issue we developed ApreciseKUre platform, which includes data on potential biomarkers, patients’ quality of life, biochemical outcomes and clinical information facilitating their integration and analysis in order to shed light on pathological characterization of every AKU patient in a typical Precision Medicine perspective [13–16].

We wanted to further elaborate and build a new database which would complement the existing ApreciseKUre database. The new database would provide the necessary underlying molecular information for novel and known clinical HGD variants. We have tried to exploit structural and sequence based information to build a predictive tool using supervised machine learning algorithm. The model has been implemented through the webserver **<u>HGDiscovery</u>**, providing functional and phenotypic consequences of HGD non-synonymous variations to better guide clinical decisions.

## Methods

### Data curation

After removal of duplicate mutations, we curated a dataset composed of 301 non-synonymous substitutions. It included 190 non-pathogenic non-synonymous variations retrieved from gnomAD v.3 (Genome build GRCh38/hg38, Ensembl gene ID: ENSG00000113924.11, Region 3:120628173-120682571) [17] and 111 AKU-causing clinical mutations. The 111 variants were first described in the study of Ascher et al. 2019 [18] and included in HGD Mutation Database (http://hgddatabase.cvtisr.sk) [19], which summarizes results of mutation analysis from approximately 530 AKU patients reported so far.

### HGD protein structure

The X-ray crystallographic 3D structure of *Homo sapiens* holo-HGD (holo-HGDHs, PDB ID: 1EY2) is incomplete; thus, it needed structural reconstruction of the missing residues of the monomer and then of the whole hexamer in order to be able to perform a complete evaluation of variants effect on protein stability and flexibility. The missing loop in the human protein structure (residues 348–355) was reconstructed by homology modeling using the *Pseudomonas putida* HGD (HGDPp) structure. By using protein BLAST [20] software we found three structures belonging to *Pseudomonas putida* with a sequence identity (the amount of characters which match exactly between two different sequences) larger than 49% and with root-mean-square deviation (RMSD) amounting to 1.8 Å for Cα [21]. We opted for HGDPp, with PDB ID 4AQ2 since, similarly to 1EY2, as it had no substrate. The structures of holo-HGDHs (PDB ID: 1EY2) and its homologous HGDPp (PDB ID: 4AQ2) were retrieved from the Protein Data Bank (PDB) [22]. Thereafter at the 1EY2 and 4AQ2 sequences alignment on BLAST web server [20], we modelled the missing residues. The modelling of the loop 348-355 was carried out using a homology model approach in which an elucidated structure of HGDPp loop was employed as template to model the structure of the protein of interest. The completed monomer structure served as a starting point for the reconstruction of the whole HGDHs oligomeric protein on the template of the asymmetric units of PDB entry 1EY2. The structure reliability was validated using PROCHECK [23]. Additionally, the energy minimization of the hexameric protein was performed using GROMACS 5.0.2 [24] in order to obtain an optimized 3D structure, a relaxation of the highly energetic conformations and a correct geometry for the following simulations (for additional information see Supplementary Methods in [18]).

### Biophysical and evolutionary score generation

A thorough structural and sequence based assessment was performed for all the HGD variants to account for the potential effects of AKU-causing mutations. Variations in protein-protein interactions between the different monomers of the hexamer HGD upon mutation was determined using mCSM-PPI2 [25]. Changes in protein stability and folding were determined using our in-house tools like mCSM-Stability [26], SDM [27] and DUET [28] and conformational flexibility changes using the normal mode analysis tool called DynaMut [29]. Effects of mutations on binding affinity of HGD to its substrate homogentisic acid were analyzed using mCSM-Lig [30]. All these are novel machine learning approaches that use graph-based signatures to represent the structural and biochemical environment of the wild-type 3D structure of a protein to quantitatively predict the effects of point mutation. To complement the above methods we used sequence based feature like SNAP2 (Screening for Non-Acceptable Polymorphisms) [31], ConSurf [32] and Provean (Protein Variation Effect Analyzer) [33] which provides valuable evolutionary information. To enrich the analysis we included protein’s wild type structural information such as residue depth, dihedral angles of the HGD chain φ (phi) and ψ (psi), relative solvent accessibility and secondary structure information. We calculated changes in molecular interactions such as hydrophobic, ionic, van der Waals’, halogen and hydrogen bonds and π interactions (cation–π, donor–π, halogen–π, carbon–π, π–π) between the wild type and mutant structures using Arpeggio [34]. We also included population-based variability using the missense tolerance ratio (MTR) [35] scoring system.

**Figure 1:**
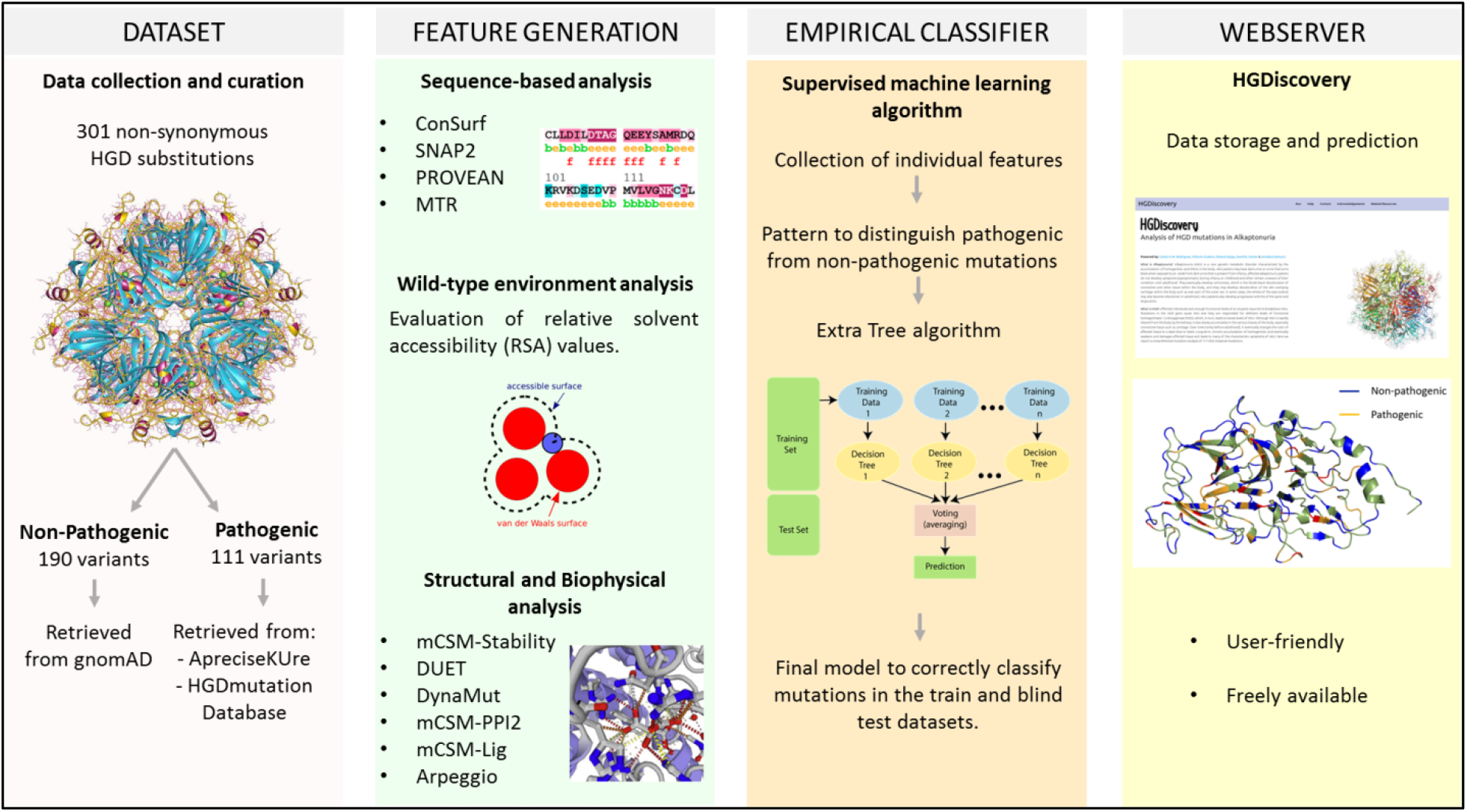
HGDiscovery workflow. The first step involves scoping published literature and clinical databases to prepare a curated list of non-synonymous HGD mutations. The second step involves generating various structure and sequence-based features for the curated missense mutations. In the third step, we use these features in a supervised machine learning algorithm to build a binary classifier, which can distinguish between pathogenic and non-pathogenic missense mutations. Finally, we develop a free available user-friendly webserver which contains phenotypic information on all HGD variants.

**Figure 2:**
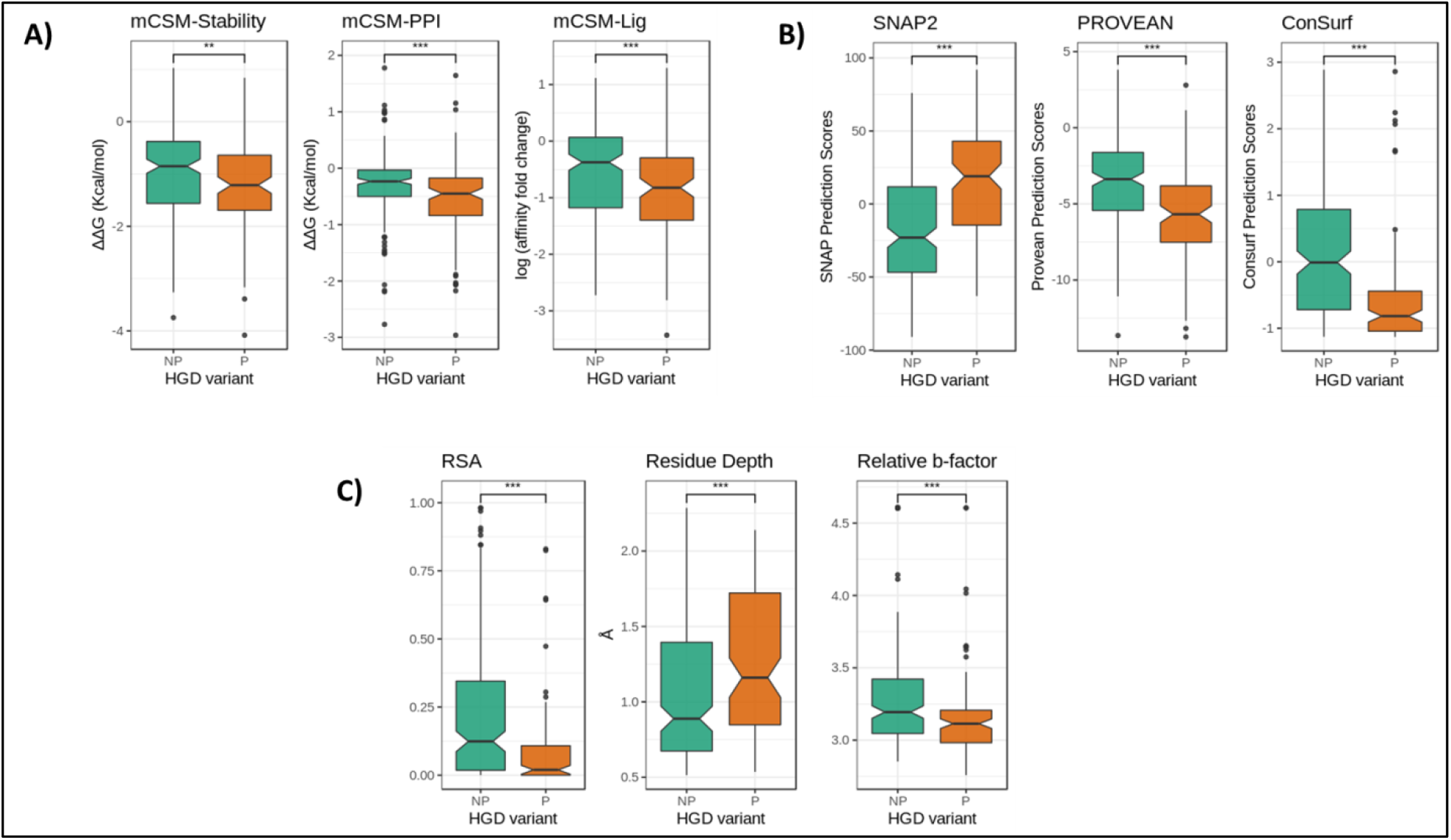
Boxplot representation of features. A) Structural features. B) Sequence-based features. C) Wild-type environment features. The non-pathogenic mutations (NP) are represented as sea green and pathogenic mutations (P) as dark orange. (*** p < 0.0001, ** p < 0.001, Welch two sample t-test).

### Supervised Machine learning for empirical model building

We evaluated different supervised machine learning algorithms for classification which is available within the scikit-learn Python library. These include – K-Nearest Neighbors (KNN), Random Forest, Decision Trees, Extra Trees, AdaBoost, Gradient Boosting, SVM, Gaussian Naïve Bayes, and Stochastic Gradient Descent. The best performing model was chosen by assessing metrics like Matthews correlation co-efficient (MCC), Receiver Operating Characteristic (AUROC) curve, accuracy, F1-score and precision. The model was trained using stratified 10-fold cross validation. We carefully split the train and blind test dataset non-redundantly with respect to the amino acid residue position.

To address the issue of imbalance between the pathogenic and non-pathogenic mutations in the data, we evaluated the model performance by both under-sampling the non-pathogenic mutations and oversampling pathogenic mutations in the train dataset [36]. The performance was compared for above mentioned scenario and the normal dataset and best results were obtained when the pathogenic mutations were oversampled using the Extra Tree algorithm. **Ex**tremely **ra**ndomized **tree** classifier (or Extra Tree) is an ensemble machine learning algorithm and a variation of the random forest algorithm. The empirical binary classifier built using this algorithm highlights a set of structural and evolutionary features which can be used to discriminate between AKU-causing and non-pathogenic variations.

### Webserver development

We have implemented HGDiscovery as a user-friendly and freely available webserver (http://biosig.unimelb.edu.au/hgdiscovery/). The front-end of the server was developed using Materializecss framework version 1.0.0, while the back-end was built in Python using the Flask framework version 1.0.2. The server is hosted on a Linux server running Apache 2.

## Results

In this work we have used the 3D protein structure to understand the functional and molecular consequences of mutations in HGD leading to AKU disease and using the information generated from these analyses we have trained a supervised machine learning algorithm to develop a predictive tool to determine novel variants which could lead to AKU manifestation. Figure 1 depicts the novel methodological pipeline we have developed.

### Sequence-based analysis of HGD variants

ConSurf, SNAP2 and PROVEAN are sequence-based predictors and consider evolutionary information to predict functionally important non-synonymous mutation. The prediction helps us understand the biological impact of a mutation on the protein structure. A consistent pattern was observed from all of the sequence based features. The pathogenic mutations were associated with deleterious scores and the non-pathogenic mutations scored neutral. All the features were statistically significant to be used to train the predictive algorithm to build the empirical tool (p-values SNAP2: 4.6 e^-14^, PROVEAN: 1.1 e^-9^, ConSurf: 2.4 e^-10^). Population-based variability was considered using the missense tolerance ratio (MTR) scoring system. Majority of the pathogenic mutations were in the bottom 25^th^ percentile, reflecting intolerance and hence associated with altering protein function.

### Wild-type environment analysis

The wild-type environment analysis includes data on relative solvent accessibility (RSA), residue depth, dihedral angles and secondary structure information for both pathogenic and non-pathogenic variants. Looking into the relative solvent accessibility values for the pathogenic and non-pathogenic mutations (p-value: 2.2 e^-8^), we see pathogenic mutations tend to be more exposed than non-pathogenic variants. It has been previously described that the HGD protomer structure constitutes of a pore in which the side chains of large number of residues are exposed [21]. These residues are thought to play an important part in the complex HGD catalytic function and we see subtle changes in the side chains as non-synonymous substitution can affect the active site functionality [18]. The residue depth values reveal pathogenic mutations are more buried than non-pathogenic mutations. This observation is congruous with earlier observation where point mutations on the surface were better tolerated in the globular hexameric HGD protein structure.

### Structural and Biophysical analysis

Our in-house biophysical tools mCSM-Stability [26], DUET [28] and DynaMut [29] were used to study and understand the impact of missense mutations on protein stability, folding and conformational flexibility. These tools are novel machine-learning algorithms which rely on graph-based signatures to calculate changes in Gibb’s free energy upon non-synonymous mutations. We observed pathogenic mutations to be associated with highly destabilizing scores affecting protein stability and dynamics. The effects of mutation on the substrate binding affinity to active site were determined using mCSM-Lig [30]. Pathogenic mutations altered the active / substrate binding pocket. mCSM-PPI2 [25] was used to assess changes in protein-protein interaction and we observed pathogenic mutations hindered the formation of the symmetrical homohexamer. Therefore, pathogenic mutations either reduced or disrupted the HGD protein activity.

### Supervised machine learning algorithm: Extra Tree

Our features could be grouped into eight distinct categories – protein stability, protein-protein interactions, ligand affinity, evolutionary conservation scores, distance parameters, MTR scores, molecular interaction and backbone geometry. Each category of features was initially used to build and evaluate the performance of the predictive model. After a thorough analysis of the individual features, we combined them together to see if there is a pattern which could be used to distinguish pathogenic from non-pathogenic HGD mutations. We observed that when different categories of features were combined together, in addition to using stratified 10-fold cross validation with Extra Tree algorithm, yielded a more robust and balanced performance. The Extra Tree algorithm implements a meta estimator that fits randomized decision trees on various sub-samples of the dataset and uses averaging to improve the predictive accuracy and reduces over-fitting [37].

**Figure 3:**
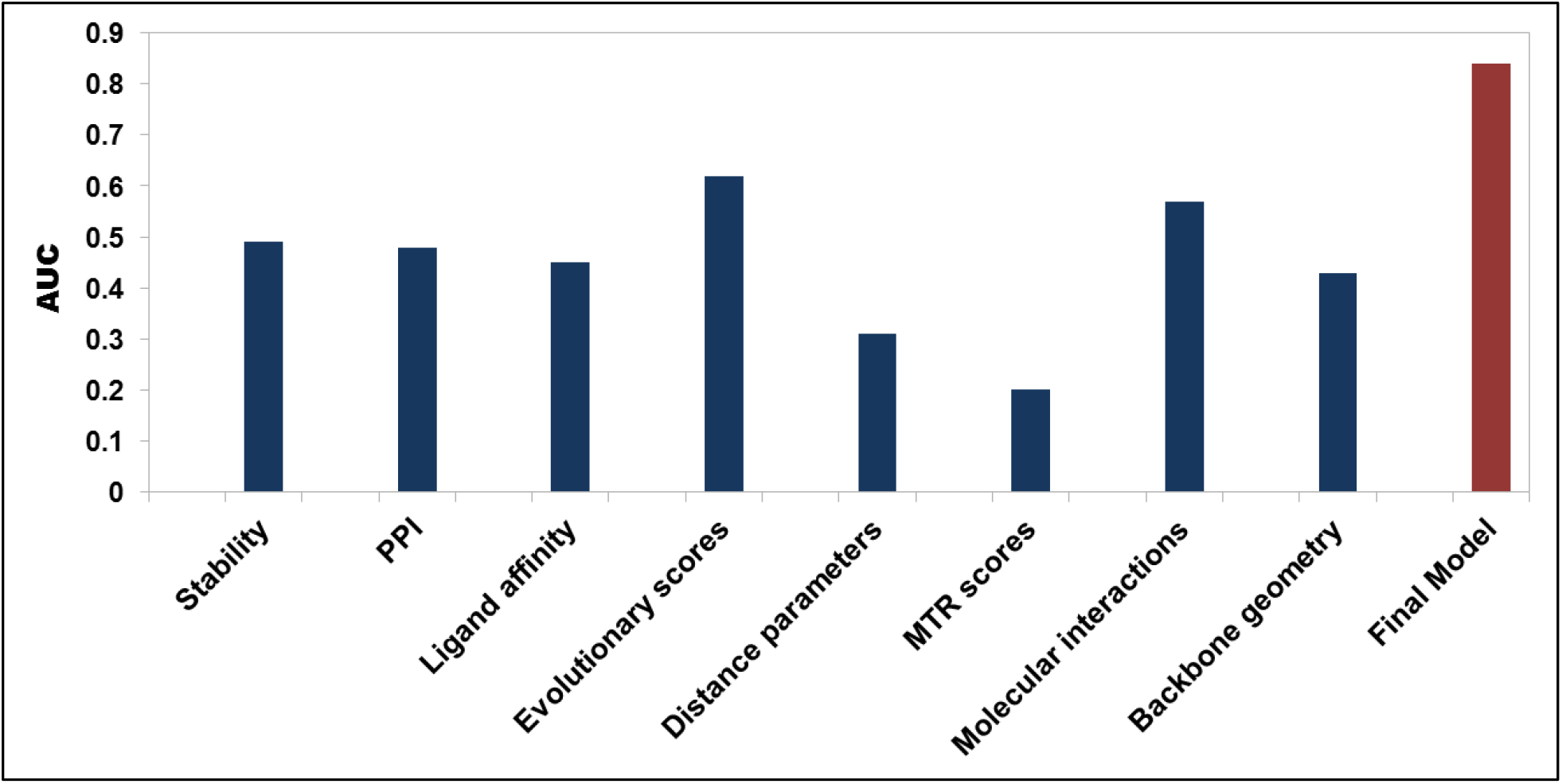
Empirical model performance trained on individual class of features. The Extra Tree algorithm was trained using stratified 10-fold cross validation using eight distinct class of features (first eight bars from left to right; dark blue bars) and with a combination of all features (red bar). The AUC score is low when a single class of feature is used for training the binary classifier, however, a significant improvement is noticed when all the eight different features are combined to build the model.

190 non-pathogenic and 111 pathogenic mutations were split into non-redundant train and blind test datasets with respect to their amino acid position. Initially we observed poor performance on the model’s ability to predict pathogenic mutation. We concluded that the train data set was imbalanced as there were more non-pathogenic mutations than pathogenic mutations. We improved the metric scores by oversampling (duplicating) [36] the pathogenic mutations in the train dataset. The final model correctly classified 84% and 73% of mutations in the train and blind test datasets respectively.

**Figure 4:**
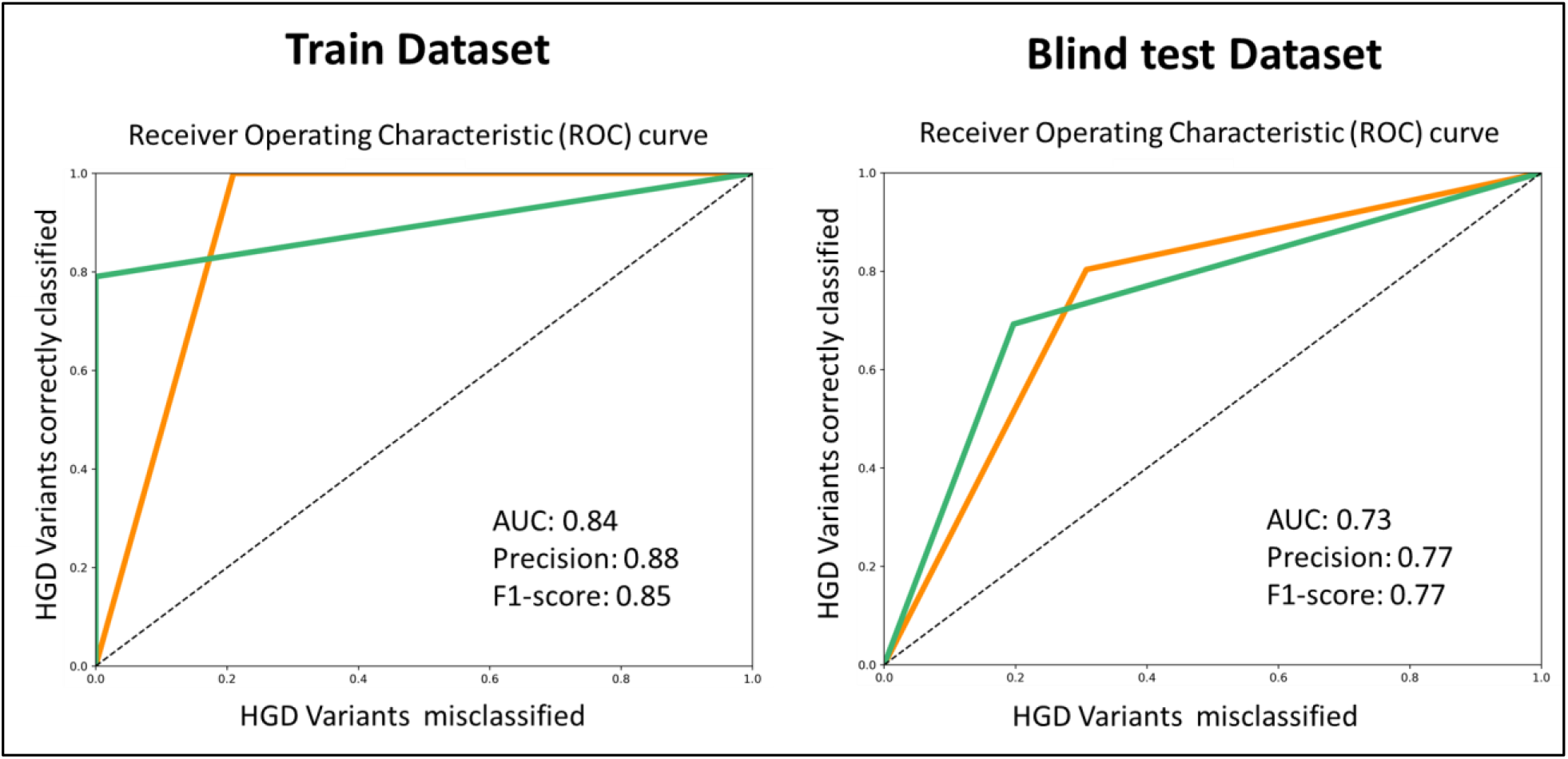
Receiver Operating Characteristic (ROC) curves of HGD classifier. The evaluation metrics shown for train and test dataset where pathogenic mutations are represented in dark orange and non-pathogenic mutations in sea green. (AUC = area under the curve).

### HGDiscovery Webserver

HGDiscovery allows for users to query for a single point mutation or submit a list of mutations to be analysed in batch. For the “Single Mutation” option users are asked to provide the point mutation as a string containing the wild-type reside one-letter code, its corresponding residue number and the mutant residue one-letter code. The “Mutation List” option requires that a text file is submitted with the list of mutations (one per line). The results page for the “Single Mutation” option displays the predicted outcome on the top alongside with details of the input mutation, wild-type residue environment, the variables and scores used by our predictive model and external links to experimental evidence (when available). An interactive 3D viewer using the NGL-viewer [38] shows the molecular contacts generated by Arpeggio [34] for wild-type and mutant structures.

On the “Mutation List” option, the results are displayed as a downloadable table. Individual analysis for each variant on the table can be analysed similarly to “Single Mutation” option by clicking the “Details” button. An interactive viewer is also shown at the bottom of the page highlighting Pathogenic and Non-pathogenic mutations on the 3D structure.

## Discussion

Here we present an empirical classifier HGDiscovery, which has phenotypic information on all variants of homogentisate 1,2 dioxygenase, (EC 1.13.11.5), an enzyme involved in the metabolism of tyrosine, whose deficiency leads to Alkaptonuria [OMIM 203500]. We combine structural, evolutionary and molecular information from known HGD variations and look to investigate a pattern to distinguish non-pathogenic from AKU-causing non-synonymous variants. So along with physiological information from ApreciseKUre platform, we have an additional AKU-dedicated database which provides new insight into functional and phenotypic consequences of novel HGD non-synonymous variations, crucial for a genetic disease like AKU to support clinical decisions.

The 3D crystal structure of the HGD active form reveals a highly complex and dynamic hexameric organization comprising two disk-like trimers [9]. An intricate network of noncovalent interactions is needed to maintain the spatial structure firstly of the protomer, the trimer and then the hexamer. This delicate structure presents a very low tolerance to mutations and can be easily disrupted mainly by missense variants compromising enzyme function. In case of HGD, missense variants represent approximately 65% of all known AKU substitutions [4, 11, 39] and 93 distinct amino acid residue positions within the structure are affected by the 111 AKU-causing missense changes. Recent studies on evolutionary conservation revealed that AKU variants were mainly located at more conserved residue positions [18] and, consequently, HGD missense changes can influence protein folding and stability or interactions with other protomers or substrate. Specifically, they can decrease stability of individual protomers, disrupt protomer–protomer interactions, or modify residues in the active-site region. Thus, when a novel HGD missense mutation is identified, it is important to distinguish causal AKU variants from non-pathogenic ones.

With sequence-based tools such as ConSurf, SNAP2 and PROVEAN we have evaluated evolutionary information in order to predict functionally important non-synonymous mutations and the biological impact of a mutation on HGD protein structure. The obtained results supported our hypothesis: the pathogenic mutations were associated with deleterious scores whereas the non-pathogenic mutations with neutral scores. Additionally, using MTR score system we have analyzed population-based variability and most of the pathogenic mutations resulted to be in the bottom 25^th^ percentile, reflecting intolerance and alteration of protein function. With the help of biophysical tools (i.e. mCSM-Stability, DUET and DynaMut) we investigated the impact of missense mutations on protein stability, folding and conformational flexibility. AKU-causing mutations appear to reduce or disrupt the HGD protein activity by destabilizing its structure and altering the active site/substrate binding pocket.

It is not uncommon that AKU patients carry compound heterozygotes for two HGD gene variants. In such cases, the estimation of the role of each missense variant is not trivial, since the hexamer could be assembled with monomers all affected by the same variant (homo-oligomer) or by two different ones (heterooligomer) [40]. Variants affecting two different regions could have additive destructive effect, on the contrary, the effects could partially compensate for those that belong to the same region. However, we do not have any tools able to evaluate such events so far [12]. Compound heterozygosity could have even interfered with our analysis, where a variant labelled as non-pathogenic could actually be pathogenic. This was the limitation of our study. But with increasing availability of genomic and clinical data after patient analysis in future, we can always update our tool and re-label the mislabeled non-synonymous variants.

The information available from the above study can be used to develop new treatment strategies, for example, use of small molecules. We know that a pathogenic mutation with destabilizing scores for stability and flexibility leading to reduced enzyme activity can be rescued partially or totally with the help of a small molecule and hence might decrease the severity of the disease [18]. Moreover, understanding the protein structure and function would also help in designing tailored drugs and therapies.

Therefore, this framework may represent an online tool that can be turned into a best practice model for Rare Diseases. We believe this is not limited to the study of AKU, but it represents a proof of principle study that could be applied to other rare diseases, allowing data management, analysis and interpretation. We applied this novel methodological pipeline to understand and determine novel drug resistant mutations in tuberculosis [41, 42] and even performed a real-time analysis [43] on tuberculosis patient. Hence, HGDiscovery is a user friendly freely available tool which could help with faster and more accurate diagnosis of AKU.

## Acknowledgements

M.K and C.H.M.R were funded by Melbourne Research Scholarships. D.B.A. was funded by a Newton Fund RCUK-CONFAP Grant awarded by The Medical Research Council and Fundacao de Amparo a Pesquisa do Estado de Minas Gerais(FAPEMIG) [MR/M026302/1]; the Jack Brockhoff Founda-tion [JBF 4186, 2016]; and an Investigator Grant from the National Health and Medical Research Council of Australia[GNT1174405]. Supported in part by the Victorian Government’s OIS Program.

## Notes

http://biosig.unimelb.edu.au/hgdiscovery/

### Competing Interest Statement

The authors have declared no competing interest.

